# Sparse Sonomyography-based Estimation of Isometric Force: A Comparison of Methods and Features

**DOI:** 10.1101/2021.08.04.455036

**Authors:** Anne Tryphosa Kamatham, Meena Alzamani, Allison Dockum, Siddhartha Sikdar, Biswarup Mukherjee

## Abstract

Noninvasive methods for estimation of joint and muscle forces have widespread clinical and research applications. Surface electromyography or sEMG provides a measure of the neural activation of muscles which can be used to estimate the force produced by the muscle. However, sEMG based measures of force suffer from poor signal-to-noise ratio and limited spatiotemporal specificity. In this paper, we propose an ultrasound imaging or sonomyography-based approach for estimating continuous isometric force from a sparse set of ultrasound scanlines. Our approach isolates anatomically relevant features from A-mode ultrasound signals, greatly reducing the dimensionality of the feature space and the computational complexity involved in traditional ultrasound-based methods. We evaluate the performance of four regression methodologies for force prediction using the reduced feature set. We also evaluate the feasibility of a practical wearable sonomyography-based system by simulating the effect of transducer placement and varying the number of transducers used in force prediction. Our results demonstrate that Gaussian process regression models outperform other regression methods in predicting continuous force levels from just four equispaced transducers and are tolerant to speckle noise. These findings will aid in the design of wearable sonomyography-based force prediction systems with robust, computationally efficient operation.

## I. Introduction

Estimation of muscle force is essential for biomechanical modelling and intuitive human-machine interaction with applications in prosthetic control and rehabilitation robotics. Surface electromyography (sEMG) correlates with muscle force and is widely used to estimate muscle forces with the help of musculoskeletal models and machine learning approaches. Typically, the mean absolute value or the peak value of the electromyography signal over a pre-defined sliding window is computed as a measure of the isometric force [1]. The derived signal is used as an activation function to drive musculoskeletal models that mathematically compute joint moments and even individual muscle forces [2]. In prosthetic control applications, machine-learning algorithms such as linear discriminant analysis (LDA) [3], support vector regression (SVR) [4], artificial neural networks (ANN) and linear regression methods [5], [6] have been extensively used for force prediction. These techniques extract time domain, frequency domain, auto-regressive features or a combination of them from the sEMG signals in order to model the force levels. Surface EMG is a non invasive modality requiring low-cost instrumentation which makes it a popular choice for several applications. However, sEMG signals suffer from poor signal to noise ratio (SNR) with dry coupled electrodes and therefore, lack the ability to finely resolve graded force levels [7]. Furthermore, cross-talk and spatial averaging in surface electrodes limits the spatiotemporal specificity of sEMG signals required for detecting activity in deep-seated muscles. High-density EMG electrode arrays [8] and invasive intramuscular EMG electrodes [9] are used to mitigate issues inherent in sEMG. Hence, there continues to be a need for a robust, non-invasive approach towards continuous muscle activity sensing that overcomes these shortcomings of sEMG based methods.

In recent years sonomyography or ultrasound imaging-based sensing of muscle activity has evolved as an attractive alternative to sEMG. Ultrasound imaging provides depth resolved information on the dynamic state of deep-seated muscle compartments in real-time. While electromyography detects the neural drive to the muscle, ultrasound senses the mechanical deformation of the muscle resulting from the motor efferent. Since, it is the muscle contraction that leads to the production of joint forces, ultrasound-based measures correlate closely to the actual output force. Several machine learning approaches have been explored to map ultrasound-derived features to joint forces or motion. Hallock et. al. have demonstrated that the cross-sectional area of a muscle directly correlates to the meausrements of isometric force at the distal joint [10]. Muscle fiber length and pennation angle have also been used as activation signals for the musculoskeletal models to estimate force from the upper and lower extremities [11], [12]. However, these studies perform computationally intensive image processing operations to extract relevant features from ultrasound B-mode sequences.This limits their utility for wearable applications where single-element transducers are used instead of transducer arrays.

Sonomyography-based control of prosthetic and other biomechatronic devices has also been widely reported in the literature. Several studies have demonstrated the potential for sonomyography to classify multiple hand gestures and individual digit movements with high degree of accuracy [13]–[16]. Proportional control of prosthetic devices by continuous tracking of upper-extremity joint angles and torques has also been demonstrated in numerous studies [16]–[19]. Although these studies established the feasibility of ultrasound-based methods as a biomechatronic control interface, most studies utilized commercial B-mode ultrasound imaging devices which limited their applicability in real-life applications. In order to address these challenges with B-mode ultrasound image-based methods, Akhlaghi et. al and Huang et. al. demonstrated that a sparse selection of ultrasound scanlines obtained from B-mode images can accurately classify multiple hand gestures [20], [21]. Similar results were also demonstrated for finger motion classification tasks with A-mode ultrasound signals obtained from wearable single-element transducers [22]–[24].

Prior studies in sonomyography-based methods for gesture classification and proportional joint force estimation utilize biomechanical changes in anatomical landmarks available in the ultrasound images by directly tracking anatomical landmarks or by tracking spatiotemporal features obtained from the ultrasound signal. Shi et. al. have demonstrated that muscle thickness [18] as well as pennation angle [25] of muscles can be tracked to provide an accurate measure of wrist angles. Other studies have focused on extracting localized spatiotemporal features such as pixel intensity changes [14], [26] from B-mode images to estimate muscle activity. On the other hand, wearable ultrasound systems employing single-element transducers utilize features such as waveform length [27], mean absolute value [24], linear regression coefficients [21], [28] etc., extracted from depth-limited windows from the A-mode ultrasound signal. However, a systematic analysis of the effect of the choice of the feature set and the modelling methodology on somomyography-based joint force or movement estimation has not been reported to the best of our knowledge.

In this paper, we propose a novel sonomyography-based method for continuous isometric force estimation from a sparse set of ultrasound scanlines. We select a small subset of ultrasound scanlines from B-mode ultrasound image sequences obtained from the forearm of healthy individuals performing ramped isometric contraction tasks. These scanlines and corresponding A-mode signals serve as analogues for single-element transducers. Our approach to feature extraction involves identifying a small set of spatial peaks in the A-mode data in order to track strongly echogenic anatomical structures. Simple spatiotemporal features are then extracted from spatial windows centered around the peaks in the A-mode signal. Thus, we extract only anatomically relevant, spatiotemporal features from the scanline derived A-mode signals greatly reducing the dimensionality of the feature space and the computational cost involved in modelling the features. We then compare the force prediction accuracy of four regression approaches - linear regression, support vector regression, ensemble bagged tree regression and Gaussian process regression. We systematically analyze the effect of the physical location of the transducer, the choice of features and ultrasound signal quality on the force prediction accuracy.

## II. Methods

### A. Subjects

We recruited five able-bodied subjects with an average age of 27±3 years, after obtaining informed consent. Subjects self-reported having no prior history of musculoskeletal disorders. Two subjects (S1 and S2) participated in the study twice, in two separate sessions conducted at least 4 months apart. Subject demographics are available in Table.I. One of the authors participated as a subject in the study. Subjects were not compensated for their involvement in the study and all study procedures were approved by the Institutional Review Board at George Mason University.

**TABLE I.** Demographics of study participants

| Subject ID | Age | Sex | Dominant arm |
| --- | --- | --- | --- |
| S1* | 30 | Male | Right |
| S2* | 27 | Male | Right |
| S3 | 23 | Male | Right |
| S4 | 24 | Male | Right |
| S5 | 31 | Male | Right |

\* Participated in two sessions at least 4 months apart

### B. Experimental setup and design

The experimental setup and data processing steps are shown in Fig.1 Subjects were seated comfortably with their dominant arm placed on a platform such that the elbow rested directly below the shoulder. The arm was secured to the platform using removable straps to restrict any movement during the experiment. A high-frequency, linear, ultrasound transducer (16HL7, Terason Inc., Burlington, MA USA) was placed on the volar aspect of the forearm approximately 4 cm to 5 cm from the olecranon process and secured with a cuff. The probe was positioned transversely, aligning the field of view to image the major flexor muscles of the forearm. The transducer was connected to a portable ultrasound imaging system (Terason uSmart 3200t, Terason Inc., Burlington MA, USA) The ultrasound system was configured to image to a depth of 4 cm with four equispaced focal zones. Image sequences from the ultrasound system were streamed to a custom developed MATLAB (version 2018b, Mathworks Inc., Natick MA, USA) interface on a laptop PC (Intel Core i7-7700HQ, 16GB RAM, 4GB NVIDIA GeForce GTX 1040Ti) in real-time using a USB-based interface (USB2DVI, Epiphan Systems Inc.). The approximate averaged frame rate obtained with the ultrasound acquisition setup was found to be 22fps.

**Fig. 1.**
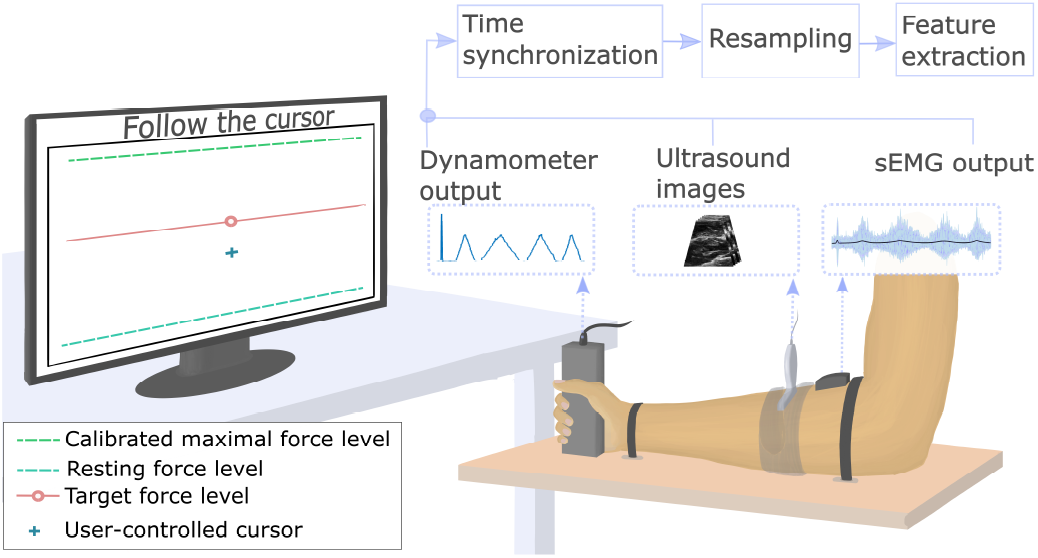
Experimental setup showing the participant instrumented with an ultrasound transducer and a single surface electromyography electrode on the forearm. The participant exerts isometric force which is measured by a hand dynamometer. The force is mapped to the movement of an on-screen cursor and the user is instructed to follow a moving target. The data obtained are synchronized, resampled and relevant features are extracted for further processing.

A wireless surface electromyography (sEMG) sensor (Trigno wireless EMG system, Delsys Inc, Boston MA, USA) was placed on the belly of the brachioradialis muscle to measure the neural activity during isometric contraction. Signals were sampled at a constant sampling rate of 1.926kHz and processed appropriately in a custom MATLAB script(see section II-C).

A USB-based hand dynamometer (HD-BTA, Vernier, Beaverton, OR, USA) was securely placed on the platform and the user was instructed to grip the dynamometer in order to measure the grip force. The dynamometer readings were sampled at 100Hz and a custom LabVIEW (Version 2020 SP1, National Instruments, Austin, TX, USA) virtual instrument was developed to read the force measurements. The VI also served as a cursor control interface for the experimental tasks described next.

#### 1) User calibration

Firstly, the user was instructed via on-screen textual prompts to produce the maximal grip force for 5 s followed by resting force of 5 s. The grip force (*F_raw_*) was measured and recorded by the VI. The resulting maximal voluntary isometric force (*F_max_*) and the resting force (*F_rest_*) values were calculated as the means of the respective 5s interval. The normalized grip force, *F_norm_* was then calculated as *F_norm_* = (*F_raw_* – *F_rest_*)/(*F_max_* – *F_rest_*). The normalized force, *F_norm_*, was therefore scaled between ‘0’ and ‘1’ and was then mapped to the vertical position of an on-screen cursor as shown in Fig. 1. The cursor moved up in response to increased grip force and moved down when the force was relaxed, settling to the bottom at resting force.

#### 2) Ramped isometric force experiments

Once the system is calibrated following the steps described above, we performed a series of experiments to simultaneously measure the ultrasound signal, surface electromyography activity and the corresponding grip force during ramped isometric contractions at varying target velocities.

Accordingly, during this part of the experimentation the user was prompted to closely follow an on-screen target (shown in red, see Fig. 1) using the cursor. At the start and end of the trials the user was instructed to produce a single, rapid twitch contraction in order to synchronize the ultrasound image sequences to the force and sEMG data. Following the synchronization twitch, the target appeared on screen and ramped up between normalized force levels of ’0’ to ’1’ and then returned to ’0’ again. Ramp sequences consisting of a ramp-up phase and a ramp-down phase lasting 20s, 30s and 40s were conducted. Three trials of each sequence were conducted in randomized order, interleaved with 10s of rest between each cycle to minimize effects of fatigue. The ultrasound image sequences, sEMG data and dynamometer force readings were processed in MATLAB as described in the section below.

### C. Data processing

The ultrasound frames, the raw EMG signal, and the dynamometer output were sampled independently on disparate software platforms as described in section II-B.1. The data had to be manually synchronized by identifying the peaks of the twitch contraction. The raw sEMG data was pre-processed by low-pass filtering with a fourth-order Butterworth filter of 800Hz cutoff frequency. Powerline interference was removed using a notch filter (center frequency = 60Hz, Q = 35). Subsequently, a moving average window of 100 ms was used for smoothing the sEMG signal. The twitch contractions at the start and end of the experiments appeared as a distinct short-duration spikes in the force data as well as the sEMG data. These peaks were identified using a peak finding algorithm and the corresponding timestamps noted for both sets of data.

In order to synchronize the ultrasound image sequences, a dummy signal was computed by obtaining the Pearson’s 2D correlation coefficient of the first image frame (assumed to be at rest) and subsequent images of the sequence. This dummy signal was then used to identify the synchronization peaks in the ultrasound image sequence.

Finally, the force data obtained from the dynomometer were downsampled to match the ultrasound sampling rate and then normalized. Nearest neighbor interpolation was used to downsample the sEMG data in order to match the ultrasound data. During data preparation, we observed that the proportion of rest state force data was considerably higher compared to the other force levels, primarily due to the design of the experiment involving interleaved resting periods. Hence, data equalization was performed to balance the data, by limiting the normalized force values between ’0.1’ and ’0.9’. Once synchronization between the ultrasound image sequences and sEMG signal were achieved, relevant features were extracted from each data and the feature extraction process is outlined in the following sections.

### D. Feature extraction

#### 1) Ultrasound features

We considered each ultrasound frame as a collection of vertical scanlines. Each scanline can be thought of as an ultrasound transducer placed on the forearm and the resultant signal is an A-mode signal from the transducer as shown in Fig. 2. Multiple scanlines can be extracted from a single ultrasound frame simulating multiple transducers placed on the forearm. Each scanline represents a depth-resolved intensity map of the underlying anatomy wherein peaks represent highly echogenic features such as bones, myofascial interfaces etc. In order to track these features, we first extracted upto four peaks in each scanline and placed a window of fixed width centered around the peaks.Thus, distinct peaks, at least 2mm apart were identified peaks with the highest intensity values were retained. At each peak window we extracted the following features:

1. Peak intensity: The peak amplitude within the window represents the echogenicity of the interface being tracked and provides a valuable measure of the importance of the feature.
2. Peak locations: In an A-mode signal, the location of the peaks represent the depth at which the anatomical features are located. Hence, the locations of the peaks identified in the previous step was chosen as a feature for our analysis.
3. Window mean: The mean value within each window centered around the peak location was calculated and considered as a feature. The mean value represents the total acoustic energy reflected by the anatomical feature within the window.
4. Window standard deviation: The standard deviation within the window of fixed width centered at each peak location was considered as a feature. The standard deviation of the window is thought to represent the sharpness of the peak and in turn, the thickness of the anatomical structure.

**Fig. 2.**
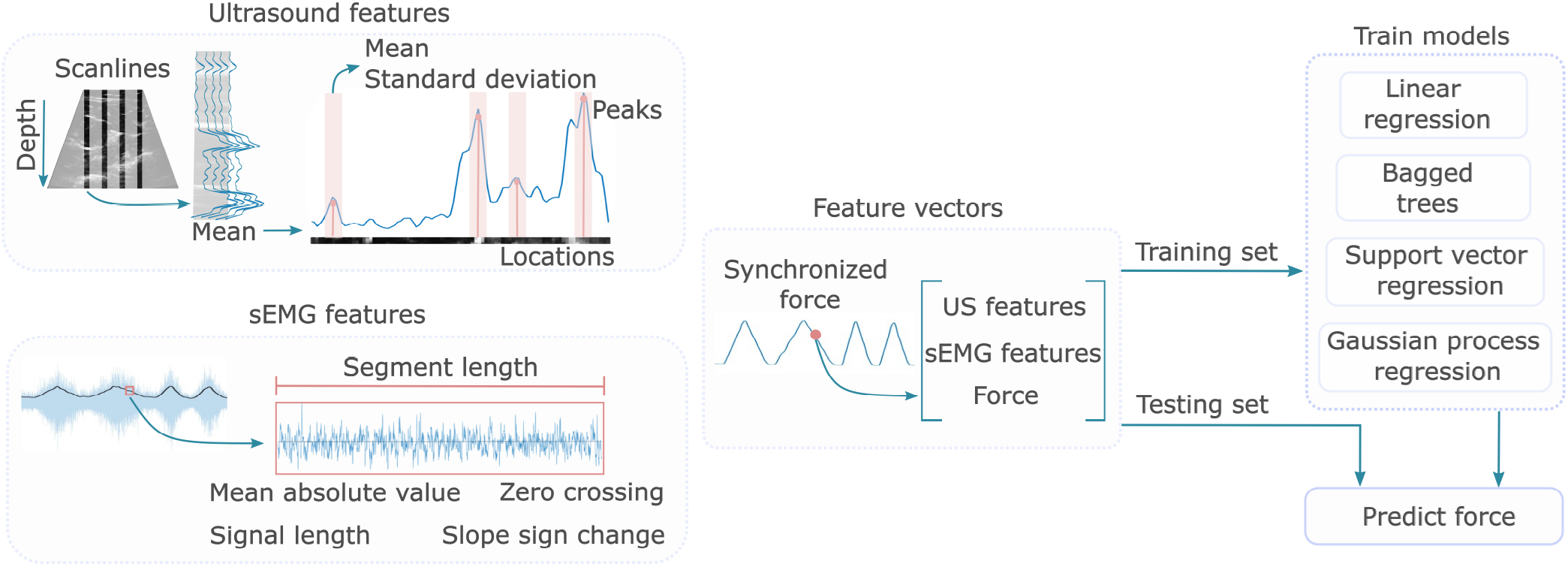
Conceptual diagram showing the process of feature extraction from ultrasound, and surface electromyography signals. The features, along with the ground-truth force measurements are fed as training data to the regression models. A holdout dataset is then used to assess the accuracy of the models and predict force levels.

#### 2) sEMG features

In addition to the ultrasound A-mode features we also included features from the sEMG signal. The sEMG signal was chopped into non-overlapping segments of length L, where L is the ratio of the total number of samples of sEMG to the number of samples of ultrasound. For each sEMG segment, we tracked four time-domain features - mean absolute value (MAV), signal length (SL), slope sign change(SS) and the number of zero crossings (ZC) [3].

### E. Training regression models

The ultrasound and sEMG features were used to evaluate four supervised regression methods: simple linear regression (LR), ensemble bagged tree (BT), support vector regression (SVR) and Gaussian process regression (GPR).

1. Linear regression: Simple linear regression model predicts the output (*ŷ*) of given test features by computing the linear combination of input features (*x*) with an added bias. *ŷ* = *θ.x* + *θ*_0_. The weights of the features, also known as the model parameters (*θ*) are generated using the training data. For a given training set with the predictors *X* and response variable *y*, The model parameters 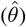 that minimize the mean square error. 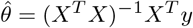 are computed.
2. Ensemble bagged trees: Bagged trees predict the output by aggregating the predictions from an ensemble of decision trees. For a given training data of size *N,* several subsets of size *n* are created using bootstrapping procedure. Bootstrapping involves creation of *m* subsets from the given training data set by randomly selecting *n* out of *N* samples with replacement. Each of these *m* subsets form a training data set for a binary decision tree. Thus, for a given input variable, *m* predictions are made. The final prediction value is the average of these *m* predictions. This reduces the variance in the predicted value. We used 20 decision trees (*m* = 20) each trained with bootstrapped data set of size same as input set. The minimum leaf size for each tress was 5.
3. Support vector regression: A linear support vector regression finds a function *f*(*x*) such that its deviation from the response variable *y* does not exceed *ε* while keeping *f*(*x*) as flat as possible.

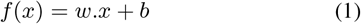 The flatness of *f*(*x*) is defined by the feature weights *w*. Therefore, the objective function requires minimization of the 2-norm of *w*, 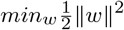. In practice, often slack variables 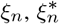 are introduced to allow outliers beyond the error bound *ε*. This modifies the objective function which tunes either the flatness of *f*(*x*) or the deviation from the tolerable error(*ε*). The modified objective function *J*(*w*) is given by,

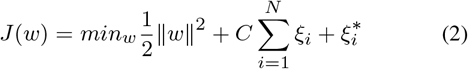

where, C is the tuning parameter. The objective function can be solved by constructing a Lagrangian dual formula by introducing non negative Lagrange multipliers *α_n_*, 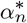 for an input *x_n_* [29], [30]. The feature weights in terms of *α_n_*, 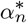 is given by,

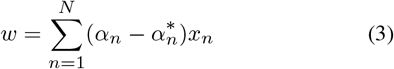 This weight parameters are used to predict the output for a test set *x_t_* as follows,

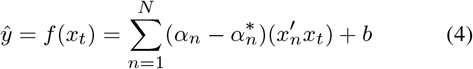 In non linear SVR, the input features are mapped to a higher dimensional space using a transformation function to improve accuracy. The non linear kernel function *G*(*x_j_,x_k_*) obtained after transformation is used in eq.4 to predict the response

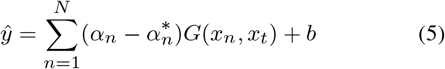 We used a polynomial SVR of order 2. The kernel function of a quadratic polynomial is given by,

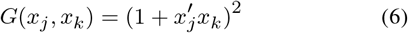
4. Gaussian process regression: In Gaussian process regression, the response *y* = {*y*_1_, *y*_2_,…*y_n_*} is assumed to be derived from a multivariate Gaussian distribution function *f*(*x*). Each function *f*(*x*) is characterized by a zero mean Gaussian distribution with covariance *k*(*x, x′*) and is derived from *x*, the set of input feature vectors {*x*_1_, *x*_2_,…*x_n_*}. *y* is expressed as,

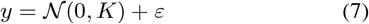

where 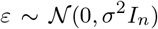 is random noise of *σ*^2^ variance and *I_n_* is an identity matrix. The covariance matrix *K*, contains the covariances *k*(*x,x′*) of given *x_i_* with every other feature vector in x and is calculated using covariance function. For rational quadratic kernel, covariance is calculated as,

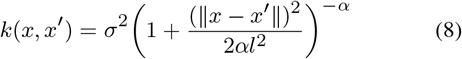

where, *α* is the scale - mixture, and *l* is the width of the kernel. To make prediction *ŷ* from the test set *x_t_*, the above estimated model parameters are used. *ŷ* is estimated as the mean value of the Gaussian distribution, *ŷ*|*y*. This is given by,

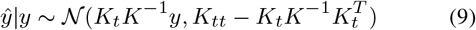
where *K_t_* is the covariance of input and test features vectors, and *K_tt_* is the covariance within test feature vectors. The predicted response is, *ŷ* = *K_t_K*^-1^*y*.

Feature vectors at each time sample were concatenated to form a composite feature matrix where each row represented an unique feature and columns represented time as shown in Fig. 2. Each feature (row) in the feature matrix was normalized. The last row of the feature matrix is the response variable or the normalized force data obtained from the dynamometer readings. 80% of the feature set were randomly selected to train the models and the remaining 20% of the data were used for testing.

### F. Analysis of factors affecting force prediction

We also investigated the effect of several experimental and measurement factors on the performance of the regression models. The specific factors considered are explained below:

#### 1) Effect of number of transducers

We analyzed the effect of the number of transducers/scanlines on the force prediction accuracy. As shown in Fig. 2, we selected equispaced columns from each ultrasound image frame to emulate the effect of placing a sparse set of single-element transducers on the forearm. In order to emulate the beam spread from a single-element transducer, we averaged five scanlines centered around the central transducer to obtain the resulting A-mode signal. We varied the number of transducers/scanlines from 2 to 10 in increments of 2. For each scanline, ultrasound features described in section II-D.1 were computed and utilised in training and testing of the regression models.

#### 2) Effect of the placement of transducers

We also analyzed the effect of the placement strategy of the ultrasound transducers on the force prediction accuracy. We compared the performance of ultrasound transducers placed at a fixed distance apart (equispaced) or at the location of maximal muscle activity. In case of equispaced transducers, equidistant scanlines were selected uniformly across the field-of-view of the ultrasound image, so as to cover the entire transverse view of the forearm, as illustrated in Fig. 2.

In order to determine areas of maximal activity in the ultrasound image sequence, we adopted the strategy reported by Sikdar et. al. [14]. The algorithm calculates an activity pattern for every subject by computing the difference between consecutive B-mode ultrasound frames. Absolute value of the image differences were then summed across time to obtain a composite activity pattern image. Higher pixel intensity in the composite activity pattern image signifies large pixel intensity changes over time. Activity scores for each scanline were obtained as the summation of intensities across the depth. Scanlines with the highest activity scores, separated by at least 1.6 mm 4 scanlines. This separation ensured that the selected scanlines were physically set apart from each other, as would be the case in a practical application.

#### 3) Effect of number of peaks and peak width in A-mode signal

We also wanted to determine the optimal number of ultrasound features and its impact on the force prediction accuracy for the regression techniques. As discussed in section II-D.1, we identified prominent peaks in the ultrasound A-mode signal from each scanline and determined simple time-domain features within these windows. We varied the number of peaks in the A-mode signal from one to four, thus varying the number of ultrasound features included in model training.

We also analyzed the effect of varying the width of the windows placed at each peak location on the prediction accuracy. The window width around the peak were varied between 2mm to 5.2mm in increments of 0.8mm. Ultrasound features were computed for each window width and utilized for training regression models. Additionally, we also calculated the width of the peak at half-maximum(Hpp) as a measure of the resolution of the anatomical structure being tracked and included it as a feature during training. The prediction results for each window width and half-maximum width were compared across the regression models.

#### 4) Effect of speckle noise in ultrasound signals

Ultrasound signals inherently contain speckle noise due to presence of subwavelength scatterers in the tissue. Speckle noise can be detrimental to image quality resulting in loss of spatial resolution [31], [32]. Therefore, we simulated the effect of multiplicative speckle noise in the ultrasound signal to study its effect on the force prediction accuracy.

The regression models were trained on clean ultrasound signals with no additional speckle noise. Speckle noise was then added to the ultrasound A-mode signals, thus degrading the signal PSNR from 60dB to 20dB. The noisy ultrasound A-mode test data were presented to the regression models and the prediction accuracy was recorded for each instance.

#### 5) Effect of adding EMG features

Finally, we evaluated the effect of combining time-domain sEMG features in addition to the ultrasound features to the regression models. The force prediction accuracy with and without sEMG features were evaluated and compared.

### G. Evaluation metrics

Our primary outcome metric to evaluate the performance of each regression model was the coefficient of determination (*R*^2^). *R*^2^ values were computed to determine the prediction accuracy of the predicted force values and the true force measurements.

### H. Statistical tests

Due to the small sample size and difficulty in verifying normality assumptions with multi-factorial parametric tests we performed Friedmann’s test to compare the performance of the regression methods. Wilcoxon’s signed-rank test was used for multiple comparisons to assess within-factor differences for the factors affecting the prediction accuracy. IBM SPSS Statistics (Version 28.0, IBM Corp, Armonk, NY, USA) was used for all statistical analysis.

## III. Results

### 1) Effect of number of transducers

Fig. 3 shows the *R*^2^ coefficient obtained for all four regression models when trained with features from a varying number of ultrasound transducers/scanlines. A significant improvement in the *R*^2^ coefficient was observed by increasing the number of transducers in all the regression methods (*p* < 0.001). Linear regression performed poorly with an average *R*^2^ of 0.764±0.17 when compared to SVR with an average *R*^2^ of 0.926±0.091. *R*^2^ of 0.978±0.01, 0.987±0.01 was noted for BT and GPR, respectively. It was observed that the mean variation in *R*^2^ when more than 4 transducers are employed is 0.001 in BT, SVR,and GPR. This was confirmed by a pairwise comparison test which showed no significant difference in *R*^2^ when number of transducers is greater than 4 for all regression methods.

**Fig. 3.**
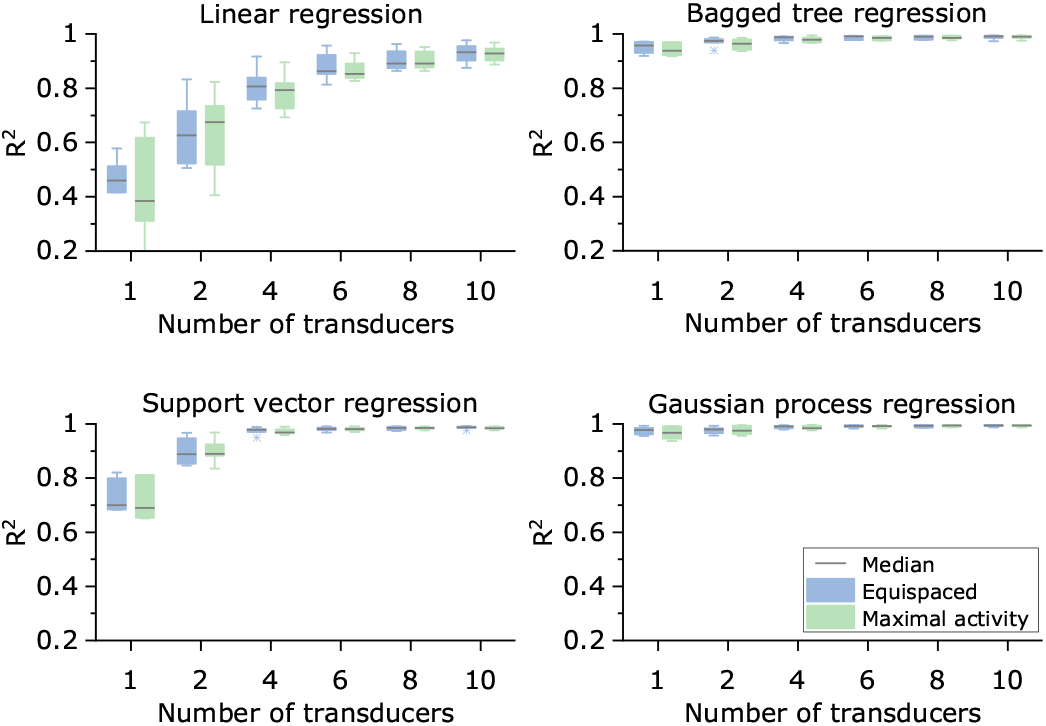
Box plots showing the effect of number of ultrasound transducers/scanlines and their placement strategy on ***R*^2^** coefficient for all regression techniques. The number of scanlines and placement strategy did not have a significant effect on prediction accuracy when four or more transducers were used.

### 2) Effect of the placement of transducers

Fig. 3 also compares the impact of the equispaced and maximal activity based transducer placement strategy on the *R*^2^ coefficient. Placement strategy had no significant impact on the force prediction accuracy in LR (*p* = 0.945), BT (*p* = 0.142), SVM (*p* = 0.812) and GPR (*p* = 0.358). Hence, based on these results, further analysis reported in the sections below were carried out with four equispaced scanlines/transducers.

### 3) Effect of using combined sEMG and ultrasound features

Fig. 4 shows the *R*^2^ coefficient when all regression models are trained with and without EMG features in addition to the ultrasound features. Pairwise comparison showed no significant difference in all methods when EMG features are added (*p* = 0.063 for LR and BT, *p* = 1 for SVR and GPR). However, the median *R*^2^ improved by 0.02 for linear regression method which was not found to be statistically significant. In view of the modest improvement offered by the addition of sEMG features particularly for linear regression models, sEMG features were included in the feature vector for results described in the following sections.

**Fig. 4.**
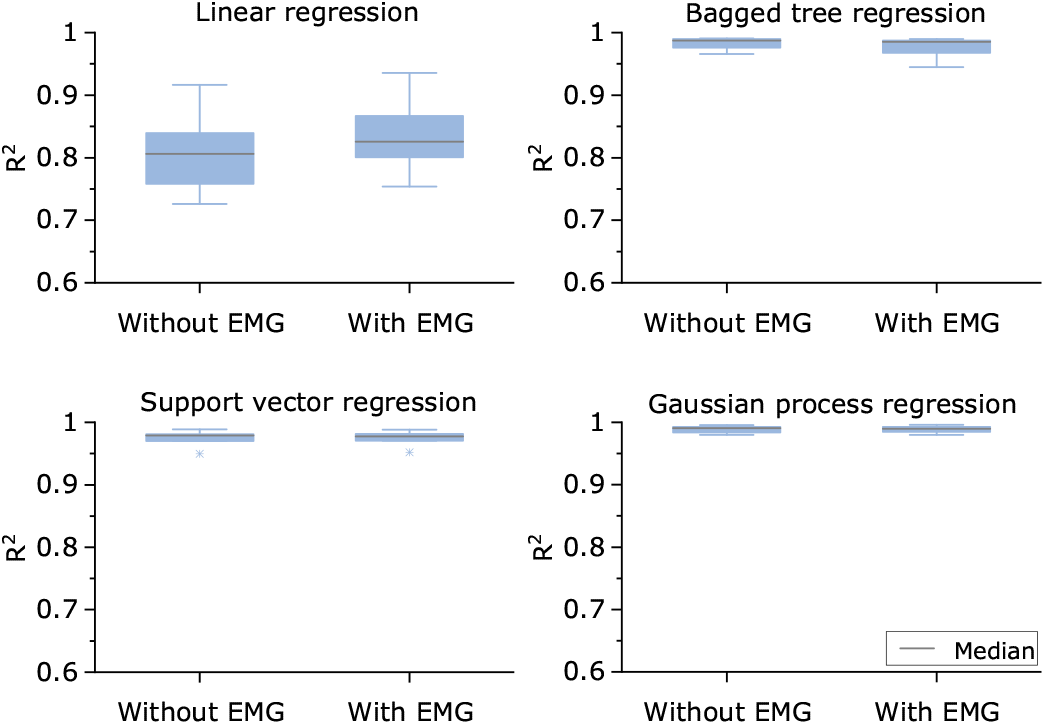
Box plots demonstrating the effect of adding electromyography (sEMG) features alongside sonomyography features on the force prediction accuracy. Addition of sEMG features did not improve the prediction accuracy significantly. Linear regression models showed a nominal increase in accuracy when sEMG features were added.

### 4) Effect of number of peaks in A-mode signal

Fig. 5 shows the effect of number of peaks on *R*^2^ coefficient for all the regression methods. LR, BT, and SVR methods showed significant improvement in *R*^2^ or force prediction accuracy as the number of the peaks included in the feature set was increased (*p* < 0.05). However, for GPR, inclusion of just one peak was sufficient to produce an *R*^2^ of 0.98±0.01 across subjects. Further addition of peaks did not lead to significant improvement in *R*^2^ for GPR (*p* = 0.867). We noticed a plateauing of the force prediction accuracy for LR, BT and SVR when three or more peaks were added to the feature set. Pairwise statistical tests confirmed that there was no significant improvement in *R*^2^ in all methods beyond the addition of three ultrasound peaks.

**Fig. 5.**
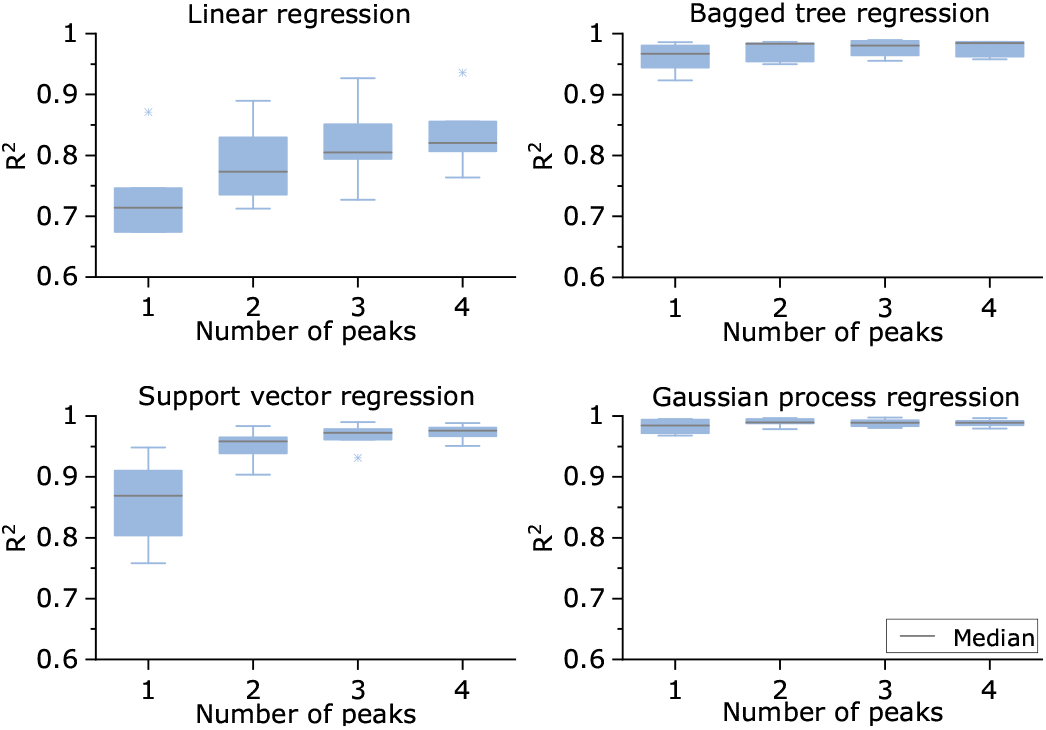
Box plots showing the effect of varying the number of peaks included as features from each ultrasound scanline or A-mode signal on the force prediction accuracy. The number of peaks had a statistically significant effect on the prediction accuracy. Gaussian procession regression model outperformed other models with ***R*^2^** of 0.98 with just one peak.

### 5) Effect of window width

We also analyzed the effect of the width of the window placed around the ultrasound peaks on the prediction accuracy and the results are shown in Fig. 6. It was found that, a window width between 2 mm and 5.2 mm did not show any significant effect on *R*^2^ in all four regression methods (*p* = 0.126 for LR, *p* = 0.137 for BT, *p* = 0.164 for SVR, and *p* = 0.321 for GPR). However, when the width was set to the full-width at half-maximum of the peak value (Hpp) and included as a feature, a significant deterioration of the prediction performance of LR (*p* = 0.004), SVR (*p* = 0.01) were observed. GPR (*p* = 0.054) and BT (*p* = 0.116) remained unaffected.

**Fig. 6.**
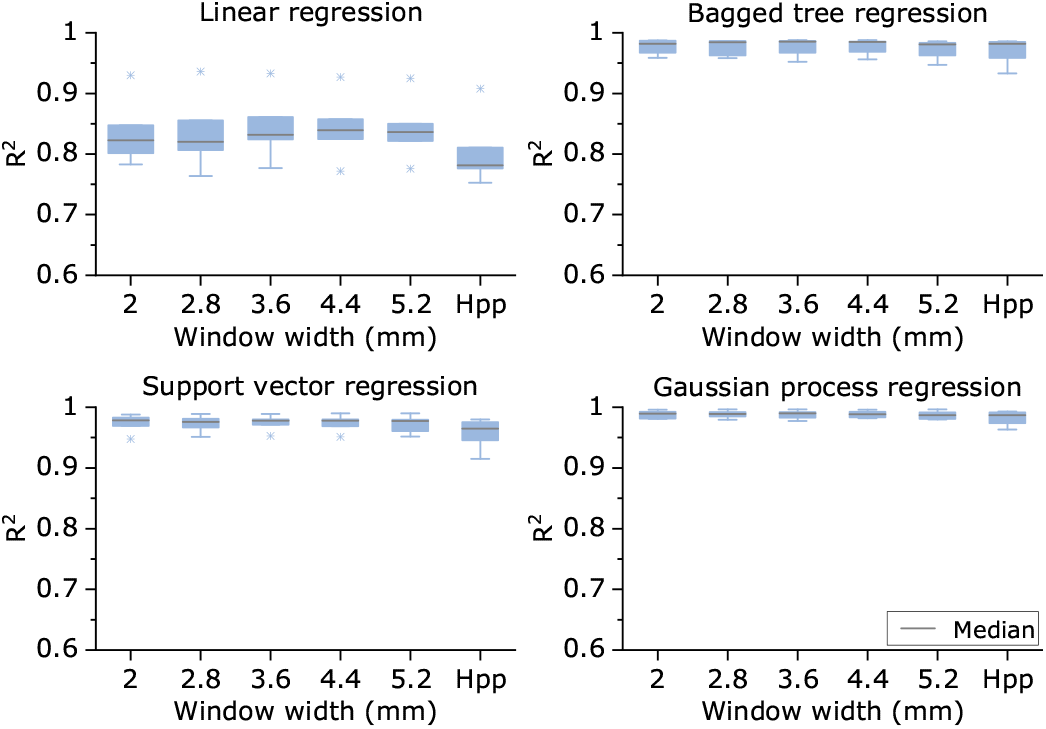
Box plots showing the effect of varying the ultrasound feature extraction window width on force prediction performance. The window width did not have a significant effect on the prediction accuracy. However, when full width at half-maximum (Hpp) was considered, accuracy deteriorated for linear regression and support vector regression models.

### 6) Effect of speckle noise in ultrasound signal

Fig.7 shows the effect of the presence of speckle noise in the ultrasound signal on the force prediction capabilities for all four regression models. It was observed that a 20 dB reduction in PSNR led to a minor reduction of prediction accuracy across all models - LR (reduced by 7%), BT (6%), SVR (10%), GPR (7%) models. However, further degradation of the ultrasound signal led to a statistically significant drop in prediction performance for all models (*p* = 0.018). When all the four models are tested with images with a PSNR of 20 dB, the *R*^2^ coefficient was found to be less than 0.5, rendering the models unsuitable for accurate force prediction.

**Fig. 7.**
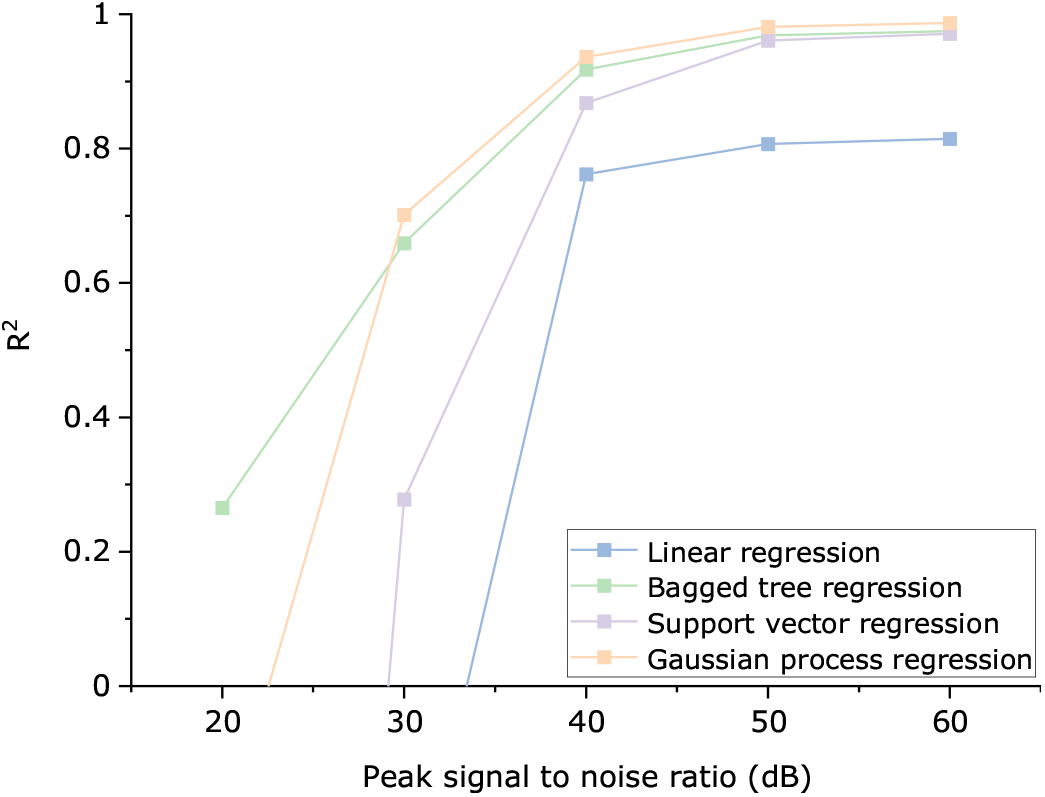
Plots showing the performance of the regression models when speckle noise was added to the ultrasound scanlines. The regression models were trained on uncorrupted data. Speckle noise was then added to the ultrasound scanlines to degrade the peak signal to noise ratio (PSNR) and the performance tested with noisy data. All models remain relatively unaffected by speckle noise levels up to 20dB.

## IV. Discussion

In this paper, we demonstrated sonomyography-based proportional isometric force estimation using four regression modelling techniques. We demonstrated that a sparse set of ultrasound scanlines consisting of a small subset of scanlines enables precise prediction of continuous force levels. Our technique utilizes simple time-domain features from a sparse set of ultrasound scanlines in conjunction with features from a single surface electromyography channel. Our approach involved computing features at prominent peak locations in the ultrasound A-mode signal. Since, the peaks in the ultrasound A-mode signal correspond to highly echogenic anatomical structures, our approach enables tracking of important anatomical features over time. It also reduces the number of features used during regression modelling by selectively tracking anatomical features of interest compared to techniques that compute features over the entire imaging depth [17], [33]. We believe that our approach will lead to increased computational efficiency and real-time prediction rates in practical sonomyographic applications.

### A. Transducer count and positioning affects force prediction accuracy

We systematically analyzed the effect of transducer sparsity on the force prediction accuracy of the regression models. Our results demonstrate that there is a significant effect of the number of ultrasound scanlines on the force prediction accuracy across all the regression methods. The performance of linear regression and support vector machine regression models improved greatly with the inclusion of features when four or more scanlines were included. However, the improvement of adding more than four scanlines/transducers were not found to be statistically significant for any technique. Our study is the first to consider and systematically analyze the trade-off between prediction performance and hardware complexity for sonomyographic proportional force prediction applications. We demonstrate that increasing the number of transducers beyond four, only marginally improves prediction performance across models, however, it also increases the complexity and cost of the hardware.

Further, we also considered the spatial placement of the transducers as an influencing factor in force prediction. We demonstrated that an equispaced sparse array of transducers outperform those placed at areas of maximal activation particularly when the number of transducers is less than four. Similar results have been reported by Akhlaghi et. al. for sonomyographic gesture recognition applications. [20]. Hence, we believe that a simple equispaced transducer array can be used in most sonomyography-based muscle activity estimation applications.

### B. EMG features marginally improve sonomyographic force prediction

We also investigated the utility of adding time-domain EMG features in addition to the ultrasound features. We found that for the linear regression model there was a small improvement in the force prediction accuracy with the inclusion of sEMG features. The other modelling approaches did not exhibit any improvement. Hence, for applications involving low transducer count using linear regression based modelling techniques, inclusion of surface electromyography features may enhance prediction performance. However, our study does not consider the effect of addition of features from sEMG transducer array co-located alongside the ultrasound transducers. Further, we do not consider frequency domain or autoregressive features which could further enhance the benefits of including sEMG features. Such a study will need specialized sensor hardware capable of simultaneous ultrasound imaging and sEMG signal acquisition.

### C. Sparse ultrasound feature set enables accurate force prediction

Ultrasound signals provide depth-resolved information on the dynamic activity of the underlying tissue. Previous approaches have attempted to exploit this information present in A-mode ultrasound signals by extracting features from fixed windows along the entire depth. This approach results in a high-dimensional feature set, requiring computationally complex approaches to model [17], [33].

On the contrary, in our approach, we identify dominant peaks in the ultrasound A-mode signal and extract features only around the identified peaks. These peaks correspond to anatomically relevant echogenic structures (such as the radius, ulna, myofascial interfaces etc.), thus limiting the information contained in the features to the motor activity only from relevant anatomical structures. We demonstrate that by considering only four such windows centered around the dominant peaks, it is possible to achieve highly accurate prediction performance.

### D. Speckle noise immune sonomyographic force prediction

Speckle is an inherent multiplicative noise present in ultrasound images which can have detrimental effects in ultrasound-based predictive models. Our results showed that even when the ultrasound signal PSNR drops by approximately 20 dB, the performance of all the models detoriorarates marginally. Further addition of speckle noise resulted in a complete failure of the linear regression and support vector regression models. The Gaussian process regression and bagged tree models were found to be more noise tolerant than linear regression and SVR models although prediction performance degraded severely. Therefore, pre-processing of ultrasound images may be necessary to remove speckle noise. However, traditional approaches towards speckle noise reduction such as median filtering and Weiner filters may not preserve anatomical boundaries that are essential for force prediction [31], [34].

### E. Limitations of the study

Our study aims to systematically analyze the various factors affecting force prediction based on sonomyography and electromyography-based techniques. The results presented in this study are based on a limited sample size. Although the models were trained to be subject-specific, inclusion of subjects from a wider age-group and gender demographic could make the results generalizable. Further, our results simulate the effect of placing single-element transducers by selecting specific scanlines from B-mode ultrasound images. This approach has been previously used in sonomyography-based gesture recognition by Akhlaghi et. al. [20] and the results have been corroborated by several studies utilizing wearable ultrasound systems [27], [28]. We intend to confirm our findings with a custom-developed wearable ultrasound imaging system in the future.

## V. Conclusion

In conclusion, we demonstrate that a sparse set of ultrasound scanlines utilizing only anatomically relevant reduced feature set can be used to predict isometric forces by regression modelling methods such as Gaussian process regression. The models are robust to a high degree of speckle noise in the ultrasound signals. The performance of the predictive models are dependent on the number of channels of ultrasound scanlines and the number of anatomically relevant features. The results of this study will help in the design of wearable sonomyography-based muscle activity detection systems.

## References

[1] Y. Na, C. Choi, H. D. Lee, and J. Kim, “A study on estimation of joint force through isometric index finger abduction with the help of sEMG peaks for biomedical applications,” IEEE Transactions on Cybernetics, vol. 46, no. 1, pp. 2–8, 2016.

[2] A. Zonnino and F. Sergi, “Model-based estimation of individual muscle force based on measurements of muscle activity in forearm muscles during isometric tasks,” IEEE Transactions on Biomedical Engineering, vol. 67, no. 1, pp. 134–145, 2020.

[3] K. Englehart and B. Hudgins, “A robust, real-time control scheme for multifunction myoelectric control,” IEEE Transactions on Biomedical Engineering, vol. 50, no. 7, pp. 848–854, 2003.

[4] H. Chen, R. Tong, M. Chen, Y. Fang, and H. Liu, “A hybrid cnn-SVM classifier for hand gesture recognition with surface EMG signals,” International Conference on Machine Learning and Cybernetics, pp. 619–624, 2018.

[5] A. Ameri, E. J. Scheme, K. B. Englehart, and P. A. Parker, “Bagged regression trees for simultaneous myoelectric force estimation,” 22nd Iranian Conference on Electrical Engineering, ICEE 2014, pp. 2000–2003, 2014.

[6] C. Choi, S. Kwon, W. Park, H.-D. Lee, and J. Kim, “Real-time pinch force estimation by surface electromyography using an artificial neural network,” Medical Engineering and Physics, vol. 32, no. 5, pp. 429–436, 2010.

[7] E. Clancy, E. Morin, and R. Merletti, “Sampling, noise-reduction and amplitude estimation issues in surface electromyography,” Journal of Electromyography and Kinesiology, vol. 12, no. 1, pp. 1–16, 2002.

[8] N. Celadon, S. Došen, I. Binder, P. Ariano, and D. Farina, “Proportional estimation of finger movements from high-density surface electromyography,” Journal of NeuroEngineering and Rehabilitation, vol. 13, 2016.

[9] E. N. Kamavuako, K. B. Englehart, W. Jensen, and D. Farina, “Simultaneous and proportional force estimation in multiple degrees of freedom from intramuscular EMG,” IEEE Transactions on Biomedical Engineering, vol. 59, no. 7, pp. 1804–1807, 2012.

[10] L. A. Hallock, A. Velu, A. Schwartz, and R. Bajcsy, “Muscle deformation correlates with output force during isometric contraction,” Proceedings of the IEEE RAS and EMBS International Conference on Biomedical Robotics and Biomechatronics, pp. 1188–1195, 2020.

[11] Q. Zhang, A. Iyer, K. Kim, and N. Sharma, “Evaluation of non-invasive ankle joint effort prediction methods for use in neurorehabilitation using electromyography and ultrasound imaging,” IEEE Transactions on Biomedical Engineering, vol. 68, no. 3, pp. 1044–1055, 2020.

[12] J. Shi, Y. Zheng, and Z. Yan, “The relationship between SEMG and change in pennation angle of brachialis,” Annual International Conference of the IEEE Engineering in Medicine and Biology - Proceedings, pp. 4802–4805, 2007.

[13] J. Shi, J.-Y. Guo, S.-X. Hu, and Y.-P. Zheng, “Recognition of finger flexion motion from ultrasound image: A feasibility study,” Ultrasound in Medicine and Biology, vol. 38, no. 10, pp. 1695–1704, 2012.

[14] S. Sikdar, H. Rangwala, E. B. Eastlake, I. A. Hunt, A. J. Nelson, J. Devanathan, A. Shin, and J. J. Pancrazio, “Novel method for predicting dexterous individual finger movements by imaging muscle activity using a wearable ultrasonic system,” IEEE Transactions on Neural Systems and Rehabilitation Engineering, vol. 22, no. 1, pp. 69–76, 2014.

[15] J. McIntosh, A. Marzo, M. Fraser, and C. Phillips, “Echoflex: Hand gesture recognition using ultrasound imaging,” Proceedings of the 2017 CHI Conference on Human Factors in Computing Systems, p. 1923–1934, 2017.

[16] A. S. Dhawan, B. Mukherjee, S. Patwardhan, N. Akhlaghi, G. Diao, G. Levay, R. Holley, W. M. Joiner, M. Harris-Love, and S. Sikdar, “Proprioceptive sonomyographic control: A novel method for intuitive and proportional control of multiple degrees-of-freedom for individuals with upper extremity limb loss,” Scientific Reports, vol. 9, 2019.

[17] X. Yang, J. Yan, Z. Chen, H. Ding, and H. Liu, “A proportional pattern recognition control scheme for wearable a-mode ultrasound sensing,” IEEE Transactions on Industrial Electronics, vol. 67, no. 1, pp. 800–808, 2020.

[18] J. Shi, Q. Chang, and Y.-P. P. Zheng, “Feasibility of controlling prosthetic hand using sonomyography signal in real time: Preliminary study,” Journal of Rehabilitation Research and Development, vol. 47, no. 2, pp. 87–98, 2010.

[19] C. Castellini, G. Passig, and E. Zarka, “Using ultrasound images of the forearm to predict finger positions,” IEEE Transactions on Neural Systems and Rehabilitation Engineering, vol. 20, no. 6, pp. 788–797, 2012.

[20] N. Akhlaghi, A. Dhawan, A. A. Khan, B. Mukherjee, G. Diao, C. Truong, and S. Sikdar, “Sparsity analysis of a sonomyographic muscle-computer interface,” IEEE Transactions on Biomedical Engineering, vol. 67, no. 3, pp. 688–696, 2020.

[21] Y. Huang, X. Yang, Y. Li, D. Zhou, K. He, and H. Liu, “Ultrasound-based sensing models for finger motion classification,” IEEE Journal of Biomedical and Health Informatics, vol. 22, no. 5, pp. 1395–1405, 2018.

[22] X. Chen, Y. P. Zheng, J. Y. Guo, and J. Shi, “Sonomyography (smg) control for powered prosthetic hand: A study with normal subjects,” Ultrasound in Medicine and Biology, vol. 36, no. 7, pp. 1076–1088, 2010.

[23] X. Yang, X. Sun, D. Zhou, Y. Li, and H. Liu, “Towards wearable a-mode ultrasound sensing for real-time finger motion recognition,” IEEE Transactions on Neural Systems and Rehabilitation Engineering, vol. 26, no. 6, pp. 1199–1208, 2018.

[24] A. J. Fernandes, Y. Ono, and E. Ukwatta, “Evaluation of finger flexion classification at reduced lateral spatial resolutions of ultrasound,” IEEE Access, vol. 9, pp. 24 105–24 118, 2021.

[25] J. Shi, Y.-P. Zheng, Q.-H. Huang, and X. Chen, “Continuous monitoring of sonomyography, electromyography and torque generated by normal upper arm muscles during isometric contraction: Sonomyography assessment for arm muscles,” IEEE Transactions on Biomedical Engineering, vol. 55, no. 3, pp. 1191–1198, 2008.

[26] C. Castellini and D. S. Gonzalez, “Ultrasound imaging as a human-machine interface in a realistic scenario,” IEEE International Conference on Intelligent Robots and Systems, pp. 1486–1492, 2013.

[27] P. Boyd and H. Liu, “A-mode ultrasound driven sensor fusion for hand gesture recognition,” Proceedings of the International Joint Conference on Neural Networks, pp. 1–6, 2020.

[28] X. Yang, J. Yan, and H. Liu, “Comparative analysis of wearable A-mode ultrasound and sEMG for muscle-computer interface,” IEEE Transactions on Biomedical Engineering, vol. 67, no. 9, pp. 2434–2442, 2020.

[29] A. J. Smola, B. Schölkopf, and S. Schölkopf, “A tutorial on support vector regression,” Statistics and Computing, vol. 14, pp. 199–222, 2004.

[30] M. Awad and R. Khanna, “Support vector regression,” Efficient Learning Machines, pp. 67–80, 2015.

[31] J. W. Goodman, “Some fundamental properties of speckle,” Journal of the Optical Society of America, vol. 66, no. 11, pp. 1145–1150, 1976.

[32] H. Rabbani, M. Vafadust, P. Abolmaesumi, and S. Gazor, “Speckle noise reduction of medical ultrasound images in complex wavelet domain using mixture priors,” IEEE Transactions on Biomedical Engineering, vol. 55, no. 9, pp. 2152–2160, 2008.

[33] X. Yang, Y. Zhou, and H. Liu, “Wearable ultrasound-based decoding of simultaneous wrist/hand kinematics,” IEEE Transactions on Industrial Electronics, vol. 68, no. 9, pp. 8667–8675, 2020.

[34] T. Loupas, W. N. McDicken, and P. L. Allan, “An adaptive weighted median filter for speckle suppression in medical ultrasonic images,” IEEE Transactions on Circuits and Systems, vol. 36, no. 1, pp. 129–135, 1989.

